# MoCETSE: A mixture-of-convolutional experts and transformer-based model for predicting Gram-negative bacterial secreted effectors

**DOI:** 10.1101/2025.08.06.668857

**Authors:** Hua Shi, Yihang Lin, Dachen Liu, Quan Zou

**Affiliations:** School of Opto-electronic and Communication Engineering, Xiamen University of Technology, Xiamen, Fujian, China; Yangtze Delta Region Institute (Quzhou), University of Electronic Science and Technology of China, Quzhou, Zhejiang, China; Institute of Fundamental and Frontier Sciences, University of Electronic Science and Technology of China, Chengdu, Sichuan, China

**Author notes:** Corresponding author (MT).

## Abstract

Identifying effector proteins of Gram-negative bacterial secretion systems is crucial for understanding their pathogenic mechanisms and guiding antimicrobial strategies. However, existing studies often directly rely on the outputs of protein language models for learning, which may lead to difficulties in accurately recognizing complex sequence features and long-range dependencies, thereby affecting prediction performance. In this study, we propose a deep learning model named MoCETSE to predict Gram-negative bacterial effector proteins. Specifically, MoCETSE first uses the pre-trained protein language model ESM-1b to transform raw amino acid sequences into context-aware vector representations. Then, by employing a target preprocessing network based on a mixture of convolutional experts, multiple sets of convolutional kernel “experts” process the data in parallel to separately learn local motifs and short-range dependencies as well as broader contextual information, generating more expressive sequence representations. In the transformer module, MoCETSE incorporates relative positional encoding to explicitly model the relative distances between residues, enabling the attention mechanism to precisely recognize the sequential relationships and long-range functional dependencies among amino acids, thereby achieving high-accuracy prediction of secreted effectors. MoCETSE has demonstrated outstanding predictive ability in 5-fold cross-validation and independent testing. Benchmark test shows that the performance of MoCETSE surpasses existing excellent binary and multi-class classifiers.

**Author Summary:** Gram-negative bacteria inject effector proteins into host cells via secretion systems, disrupting normal cellular functions and inducing diseases. Accurately identifying these virulent proteins is key to understanding bacterial pathogenic mechanisms and developing therapies. However, existing methods face issues like feature redundancy, inadequate capture of long-range dependent signals, and low computational efficiency. We developed MoCETSE, a novel computational method enabling end-to-end intelligent prediction of effector proteins from raw sequences. Due to the high computational cost of position-specific scoring matrix encoding, we use pre-trained protein language models to extract structural, evolutionary, and functional features from sequences, providing biologically meaningful inputs for subsequent deep learning models. Our hybrid convolutional expert network reduces dimensionality of high-dimensional embeddings and extracts multi-scale features, effectively overcoming feature redundancy and information loss, and improving model performance and efficiency. In learning secretion signal features, relative positional encoding models amino acid order, capturing critical long-range dependent signals, and enhancing the biological interpretability of predictions. MoCETSE outperforms existing tools like DeepSecE in cross-category predictions, offering a high-throughput method for effector protein prediction and clues for studying bacterial infections and developing therapies.

## Introduction

Secretion systems are complex molecular apparatuses unique to Gram-negative pathogens. They provide a unique pathogenic mechanism by mediating the transmembrane transport of effector proteins for direct injection into the host cytoplasm [1,2]. Currently identified bacterial secretion systems are mainly classified into type I (T1SS), type II (T2SS), type III (T3SS), type IV (T4SS), and type VI (T6SS) based on their secretion mechanisms and molecular characteristics [3]. Although there is heterogeneity in the effector proteins carried by different pathogens, these secretion systems play an indispensable role in the infection process of various clinically important Gram-negative bacteria by regulating the delivery of key virulence factors [4,5]. For example, pathogenic bacteria represented by *Shigella* and *Citrobacter rodentium* can inject type III secreted effectors (T3SEs) into host cells through T3SS [6], directly disrupting the normal physiological activities of host cells. The pathogenicity of *Legionella pneumophila* depends on the establishment and maintenance of *Legionella-containing* vacuoles regulated by Icm/Dot T4SS [7]; this system specifically interferes with host vesicle transport, actin reorganization, and signal transduction by transporting T4SEs into the host cytoplasm. The T6SS of *Pseudomonas aeruginosa*, as an efficient protein transport apparatus, can directly deliver type VI secreted effectors (T6SEs) across the membrane to adjacent target cells [8], enhancing its pathogenic ability during infection. Therefore, predicting, identifying, and classifying bacterial secreted effectors and understanding the virulence processes of secreted effectors are of great significance for an in-depth understanding of microbial pathogenic mechanisms and the development of new anti-infective therapeutic strategies.

To date, the published secretion substrate identification tools include type I secreted effectors (T1SEs) [9], T3SEs [10–12], T4SEs [13–17], and T6SEs [18,19]. Among them, feature encoding-based secreted protein prediction models have made significant progress. Algorithms represented by Bastion3 achieve accurate prediction of T3SEs by integrating the evolutionary conservation and structural constraint information contained in the position-specific scoring matrix (PSSM) [10]. CNN-T4SE, on the other hand, integrates three feature encoding strategies: PSSM, protein secondary structure and solvent accessibility (PSSSA), and one-hot encoding, thereby significantly improving the identification accuracy of T4SEs [14]. However, these traditional encoding strategies have high computational costs, are difficult to effectively capture the features of homologous sequence proteins, and cannot fully explore the synergistic relationship among sequence, structure, and function.

Protein language model (PLM) [20–22] have opened up a new research direction for effector prediction tasks. Such models are based on the attention mechanism [23] and mine potential biological laws from data containing billions of protein sequences through deep neural networks in large-scale unsupervised learning tasks. PLM can convert raw amino acid sequences into distributed representations with rich biological significance [24,25], thereby capturing the structure, function, sequence characteristics, and evolutionary relationships of proteins.

Existing studies have shown that applying PLM to the task of secreted substrate classification can significantly improve the accuracy and robustness of prediction models. DeepSecE, by integrating large-scale PLM with a transformer module optimized for secretion signals, demonstrates a strong capability in efficiently identifying disease-related proteins in bacterial genomes [26]. T4Seeker combines distance residue (DR) features, long short-term memory (LSTM) features, and PLM features. Compared to current T4SE identification models, it shows significantly improved prediction performance [17]. However, due to the significant heterogeneity of secretion signals in sequence length, spatial position, and combination patterns [27], traditional protein representation extraction methods have difficulty fully capturing their long-range dependencies and multimodal features. Therefore, developing new methods that can effectively extract multi-scale and multimodal features from PLM outputs is key to improving the performance of secretory system effector protein prediction models. In addition, traditional effector protein prediction methods have obvious shortcomings, as they cannot effectively capture the inherent sequential features in protein sequences, leading to the model’s significantly insufficient ability to recognize the positional correlation of conserved motifs and long-distance functional dependencies. This defect often seriously affects the model’s performance in effector protein function prediction tasks and results in a lack of interpretability of the model.

To solve these shortcomings, we first constructed a target preprocessing network (TPN), a module based on hybrid convolution experts, designed to focus on the key sequence features output by PLM. Compared with the simple 1D convolution layers adopted by existing prediction models, target preprocessing network can recognize multi-level features of proteins, thereby improving the fineness and comprehensiveness of feature extraction. Secondly, we introduced a transformer module incorporating relative position encoding. Compared with traditional absolute position encoding, the former can further enhance the model’ s perception of the relative order and functional associations between amino acid residues. Based on this, we developed a powerful effector protein prediction model, namely MoCETSE, which can accurately identify five major secreted effectors (T1SE, T2SE, T3SE, T4SE, and T6SE). This model not only rivals current advanced binary classifiers (such as Bstion3 [10], CNN-T4SE [14], and Bastion6 [18]) in performance, but its prediction accuracy also surpasses the existing popular multi-class classifier DeepSecE [26]. Through the unique feature extraction and modeling capabilities of the target preprocessing network and relative position encoding, MoCETSE provides a new research perspective for in-depth understanding of the functional roles of effector proteins in bacterial pathogenic mechanisms, and is expected to promote further development of bacterial effector protein function annotation and pathogenic mechanism research.

## Materials and methods

### Data description

The non-redundant labeled training and test sets used by the MoCETSE effector protein classifier were all derived from the multi-class effector protein prediction tool DeepSecE [26]. The training set contains 1,577 non-effector proteins, 128 T1SEs, 68 T2SEs, 406 T3SEs, 504 T4SEs, and 232 T6SEs; the test set consists of 150 non-effector proteins, 20 T1SEs, 10 T2SEs, 30 T3SEs, 30 T4SEs, and 20 T6SEs. The protein sequences in these training and test sets span a wide range of lengths, from fewer than 100 amino acids to over 1,000 amino acids. As shown in <u>S1 Fig</u>, the lengths of these proteins are mainly distributed within the range of 100–600 amino acids. During the benchmark testing phase, for type III, IV, and VI secretion proteins, we used the test datasets from Bastion3 [10], CNN-T4SE [14], and Bastion6 [18] as benchmark data, comparing the MoCETSE model with existing popular classification methods. Since there is no independent test data for type I secretion proteins, we constructed a benchmark test dataset using 20 T1SEs and 150 non-secretion proteins.

### Overview Framework of MoCETSE

Our proposed innovative framework, MoCETSE (Fig 1A), starts with the input of effector protein sequence data. We leverage ESM-1b [21], a pre-trained protein language model, to encode amino acid sequences into meaningful feature vectors. Specifically, ESM-1b is constructed by stacking 33 layers of transformer modules, which alternate between self-attention mechanisms and feed-forward connections. Each transformer module contains multiple attention heads, which can capture information in the sequence from different perspectives. We limit protein sequences to 1020 amino acids and then input them into the ESM-1b model, where they are converted into tokens, which are then output as tokens embedded with 1280 dimensions. To enhance the model’s learning ability for protein sequence features and training efficiency, a target preprocessing network (Fig 1B) is utilized. This network reduces the token embedding dimension from 1280 to 256 by leveraging eight convolutional experts that work collaboratively. Subsequently, the features are fed into a transformer module incorporating relative position encoding (Fig 1C), where the number of attention heads is 4, and the feed-forward network internally uses GELU as the activation function [28]. After processing by the transformer module, the obtained feature representations pass through a fully connected layer for linear transformation, mapping them to a dimensional space equal to the number of classification categories. Finally, the softmax activation function is used to convert the output values into a probability distribution, ultimately achieving the classification of non-effector proteins and five types of effector proteins (T1SE to T4SE and T6SE):

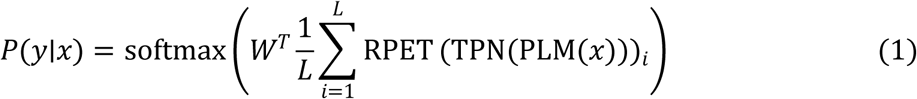

**Fig 1.**
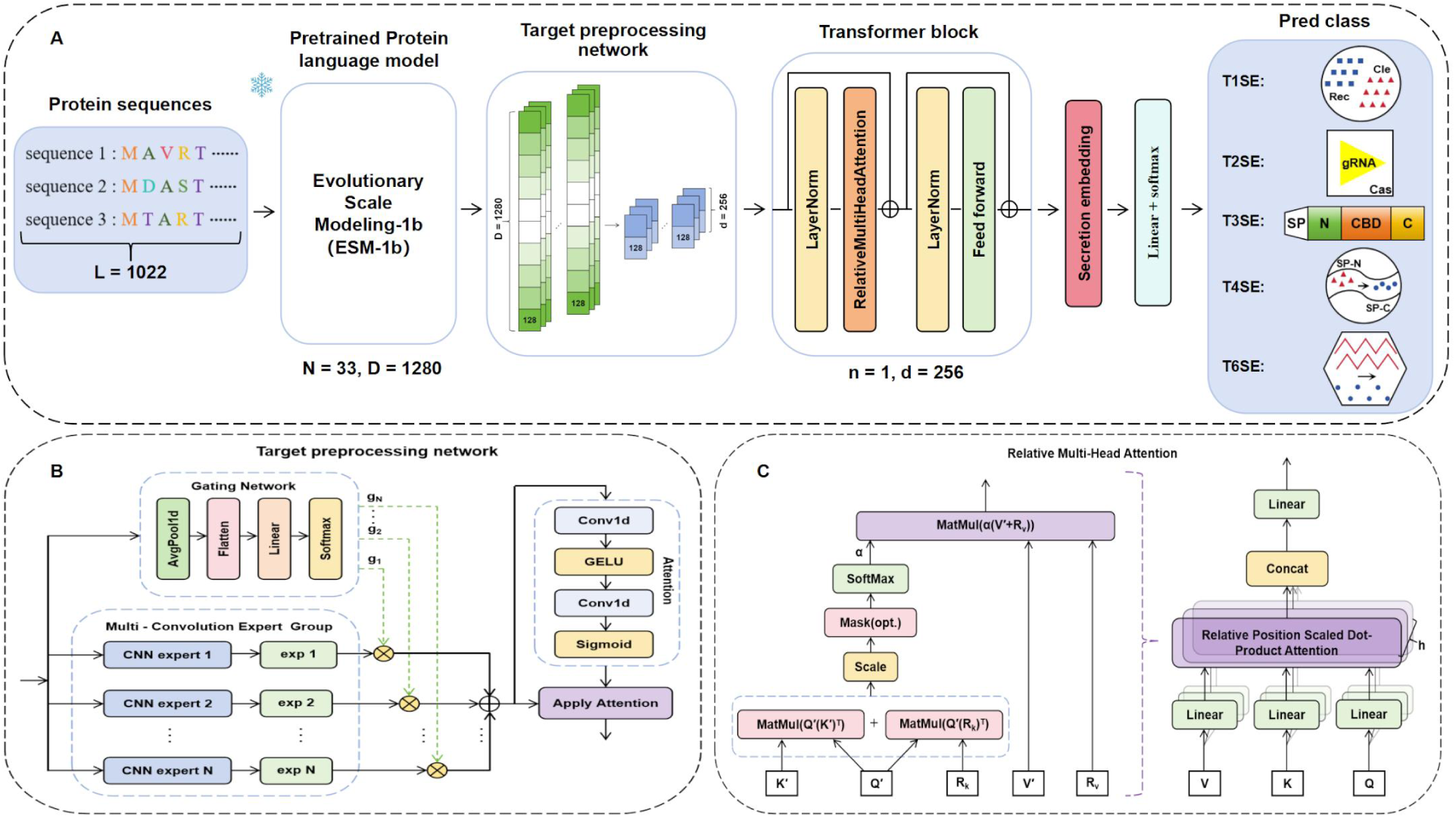
The Overall workflow of MoCETSE. (A) MoCETSE uses the pre-trained protein language model ESM-1b and a transformer module to achieve effective learning of secreted protein features. A snowflake icon on the ESM-1b module denotes frozen weights that remain unchanged during subsequent training. (B) A target preprocessing network based on a mixture-of-convolution-experts architecture is incorporated to capture a broader range of discriminative features from secreted effectors. (C) Relative position encoding is integrated into the multi-head attention mechanism, enhancing the model’s capacity to recognize key functional motifs within secreted effectors.

Where *x* denotes the input protein sequence, *L* indicates the sequence length, (.)*_i_* represents the embedding vector of the *i*th amino acid residue, PLM represents the pre-trained protein language model, TPN represents the target preprocessing network, and RPET represents the relative position-encoded transformer.

### Protein language model

Numerous studies have confirmed that the feature representations generated by protein language models can effectively capture the sequence characteristics of proteins. This advantage has led to their extensive application in predicting protein structure and functional properties [29,30]. ESM-1b employs an unsupervised learning strategy to learn and extract various biological embedding features of proteins from the UniRef50 database [31], which contains approximately 250 million protein sequences. This database systematically integrates protein sequences from the UniProt Knowledgebase using a clustering algorithm, grouping sequences with at least 50% sequence similarity into the same cluster. The initial weights of ESM-1b are obtained from the pre-trained model parameter repository (https://github.com/facebookresearch/esm) and remain frozen without any fine-tuning during subsequent training.

### Target preprocessing network

The target preprocessing network (TPN) preprocesses the high-dimensional embedding vectors generated by the protein language model, reducing the feature dimension while focusing on the key features of the sequence. Unlike conventional single-layer 1D convolutional approaches that fixedly extract features, this network employs a multi-convolution expert architecture to capture diverse features from multiple receptive fields, thereby improving the richness and flexibility of feature extraction. In addition, an integrated attention mechanism allows the model to assign greater weight to task-relevant regions while suppressing noise and irrelevant information, further enhancing the quality of the learned features.

The target preprocessing network is designed following a divide-and-conquer strategy and is implemented using a mixture of experts (MoE) architecture. Originally proposed by Jordan and Jacobs in 1991 [32], MoE is a classical machine learning framework that enables efficient modeling of complex data distributions by decomposing tasks into subproblems, assigning them to specialized expert modules, and integrating their outputs through a gating mechanism. Recent advances, such as DeepSeekMoE [33], demonstrate that fine-grained expert partitioning can achieve competitive performance with reduced parameter counts, while DynamicMoE [34] employs a dynamic gating network to adaptively activate a subset of experts based on input complexity, balancing computational cost and model accuracy. These studies highlight the effectiveness of the mixture of experts paradigm. As illustrated in Fig 1B, the MoCE-based TPN in this work comprises three key components: a gating network module [35], a multi-convolution expert group [36], and an attention module [23].

The gating network module serves as the central coordinator within the target preprocessing network, dynamically assigning weights to individual convolutional experts. By leveraging the specialized capabilities of different experts, the gating mechanism facilitates task decomposition and mitigates interference that may arise from globally shared weights in conventional single-network architectures. Specifically, it generates a set of gating weights *g*_1_, *g*_2_, ⋯, g hrough a softmax activation, producing a probability distribution that adaptively modulates the contributions of each expert. The final output is obtained by computing a weighted sum of the expert outputs, with the gating weights determining each expert’s relative influence. The output of the gating network module *g*_i_*x*_i_ is defined as follows:

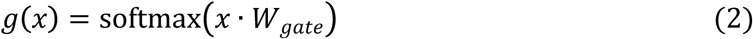

Where *x* represents the feature vector of raw input data after preprocessing via average pooling and flattening, and W*_gate_* represents the learnable weight matrix.

As a pivotal component of the target preprocessing network, the multi-convolution expert group module integrates the parallelism of the mixture of experts architecture with the feature extraction capabilities of convolutional neural networks (CNNs) to form a differential feature extraction system. This module comprises multiple parallel CNN sub-modules, each acting as an expert branch with its own parameter set, enabling it to extract features at different spatial scales and dimensions from the input data. The gating network assigns a weight to each expert branch, and these weights are used to perform a weighted fusion of the expert outputs. This design enables the module to leverage the strengths of multiple experts, adaptively integrate diverse features, and enhance the TPN’s capacity to model complex data representations effectively.

For a given input *x*, let *g_i_ x*_i_ represent the output of the gating network, and let *E*_i_ *x*_i_ represent the output of the *i*th convolutional expert network. The output *E*_expert_ of the multi-convolution expert group module can be written in the following form:

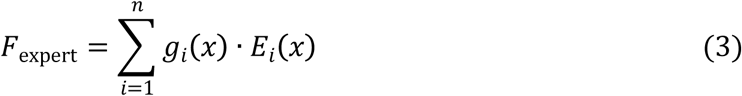

To enhance the selective emphasis on informative features, a lightweight attention module is incorporated into the target preprocessing network. This module is designed to refine and filter features by dynamically adjusting their importance. It employs the GELU activation function to improve the non-linear representation capacity and uses a sigmoid function to generate an attention weight distribution. These weights are applied to the fused feature representation, which is obtained from the interaction between the multi-convolution expert group and the gating network, through an Apply Attention mechanism. This process adaptively emphasizes task-relevant information while suppressing irrelevant or noisy features, thereby improving the network’s ability to process complex input data. The fusion feature *F*_expert_, cooperatively generated by the multi-convolution expert group and the gating network, serves as the input to the attention module, where the attention module produces an element-wise attention mask through convolutional operations, GELU activation, and sigmoid normalization. The output

*F*_output_ of the final TPN can be written in the following form:

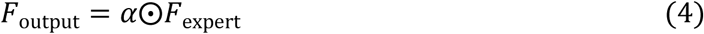

### Relative Multi-Head Attention

The transformer model proposed by Vaswani et al. in 2017 [23] relies entirely on the attention mechanism in its architectural design and has achieved breakthrough progress in the field of machine translation. It is worth noting that the transformer model does not explicitly model relative position information or absolute position information in its architecture, but instead requires additional absolute position representation information to be added to the input. In the transformer model, the attention mechanism calculates the output matrix as follows:

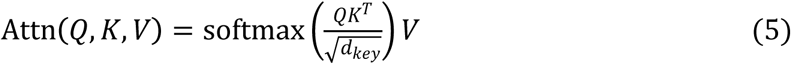

Where *Q*, *K* and *V* are the matrices integrated by the query vector, key vector, and value vector respectively, and *d_key_* represents the dimension of *K* and *V*.

Unlike recurrent neural networks [37] and convolutional neural networks [38] commonly used in sequence learning tasks, the transformer model neither relies on convolution operations nor abandons the recurrent structure, which makes it completely lack sensitivity to the inherent order of sequences. Therefore, models based on the attention mechanism usually adopt position encoding to solve this problem. As shown in Fig 1C, this study incorporates relative position encoding [39,40] into the multi-head attention mechanism of the transformer, enabling MoCETSE to more accurately capture the relative position dependence of key motifs in the sequence in the effector protein prediction task and enhance the generalization ability to sequences of different lengths. A relative position scaled dot-product attention module can be expressed as:

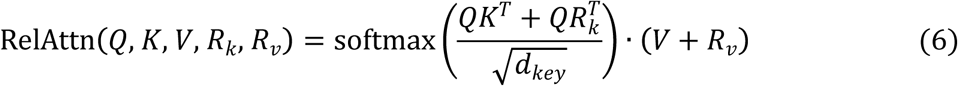

Where *R_k_* and *R_v_* represent the relative position encoding matrices for position bias and position correction, respectively.

The relative position multi-head attention mechanism incorporates relative positional encoding into the computation of attention weights. This allows the attention scores to be influenced not only by the content-based similarity between queries and keys, but also by their relative positional relationships. The output of the relative position multi-head attention mechanism is defined as follows:

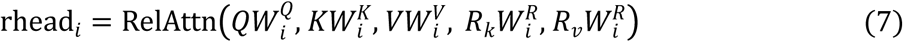

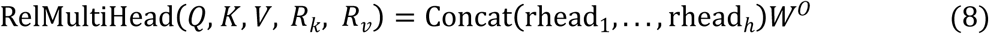

Where *W_i_^Q^*, *W_i_^K^*, *W_i_^V^*, *W_i_^R^* and *W°* related matrices represent learnable parameters. The MoCETSE model, by virtue of the multi-head attention mechanism with relative positions, precisely focuses on key regions in protein sequences (such as signal peptides at the N-terminus or C-terminus, specific motifs, etc.), thereby learning specific representations that can effectively distinguish between different types of secreted effectors and non-secreted effectors.

### Performance assessment

In both 5-fold cross-validation and independent testing, model performance was evaluated using macro-average score, F1 score, area under the receiver operating characteristic curve (AUC), and area under the precision–recall curve (AUPRC). For the task of predicting five types of secretory proteins, the receiver operating characteristic (ROC) curve provides a visual representation of the model’s ability to distinguish between positive and negative samples across different threshold settings. In addition, the multi-class confusion matrix presents a detailed view of the classification outcomes for each protein category. A higher AUC value (closer to 1) indicates better discrimination between positive and negative samples [41], while a higher AUPRC value (closer to 1) reflects stronger capability in identifying positive instances [42].To compare performance with existing secretory protein prediction and classification methods, several evaluation metrics were used, including accuracy (ACC), recall (REC), precision (PR), F1 score (F1), and Matthews correlation coefficient (MCC), providing a comprehensive assessment.

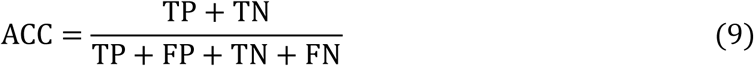

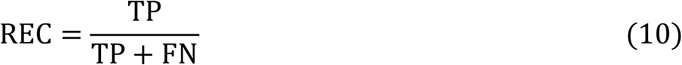

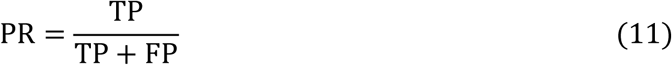

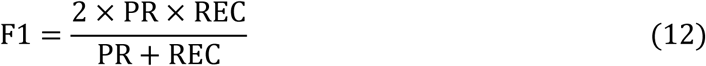

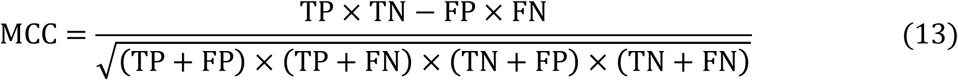

Where TP, TN, FP, and FN represent the numbers of true positives, true negatives, false positives, and false negatives, respectively. All evaluation metrics were calculated using the Scikit-learn library in Python.

### Sequence significance map

MoCETSE leverages the relative position multi-head attention mechanism within the transformer module to infer the importance of individual amino acids in a protein sequence by encoding the relative distances between residues. Specifically, the model quantifies the contribution of individual amino acids to the protein secretion process by deriving significance scores from attention matrices. This conversion involves aggregating outputs from all attention heads and incorporating relative positional information to compute a weighted average that reflects the influence of a given residue on all other positions in the sequence. The mathematical formulation is as follows:

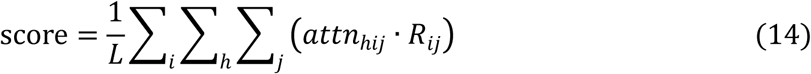

Where *R_ij_* denotes the relative positional weight between amino acids at positions *i* and *j*, and *L* represents the length of the protein sequence. The term *attn*_ℎij_ refers to the attention weight between positions *i* and *j*, as computed by the ℎth attention head. A higher value of attn hijindicates a stronger association between the two positions. To visualize the learned positional importance, the Python toolkit Logomaker [43] is used to generate the sequence significance map. This map provides an intuitive visualization of the sequence features characteristic of secretory proteins, highlighting biologically relevant properties within key regions.

## Results

### UMAP visualization of secretion protein embeddings

To evaluate the discriminative ability of the protein embedding features learned by the model, we applied the Uniform Manifold Approximation and Projection (UMAP) algorithm to perform dimensionality reduction and visualization analysis on the feature representations of the training set [44]. UMAP mapped the high-dimensional features generated by MoCETSE and the pre-trained language model ESM-1b into a two-dimensional space to intuitively present the distribution relationships among the samples (Fig 2). The results showed that the embeddings of MoCETSE formed clear clusters in the low-dimensional space: the five different types of effector proteins and non-effector proteins each aggregated into distinct clusters, with clear boundaries between them. Further observation revealed that samples within the same category were tightly clustered, indicating high consistency and homogeneity of sequence features. In contrast, the clusters of different categories displayed significant spatial separation, demonstrating that MoCETSE effectively distinguished the sequence features of varying protein types during the encoding process. This stands in sharp contrast to the projection of ESM-1b, which exhibited a greater degree of overlap between categories and less distinct clustering boundaries. This intuitive and robust clustering phenomenon indicates that MoCETSE not only captures class-specific sequence features related to the secretion system but also significantly enhances the recognition and classification ability of effector proteins.

**Fig 2.**
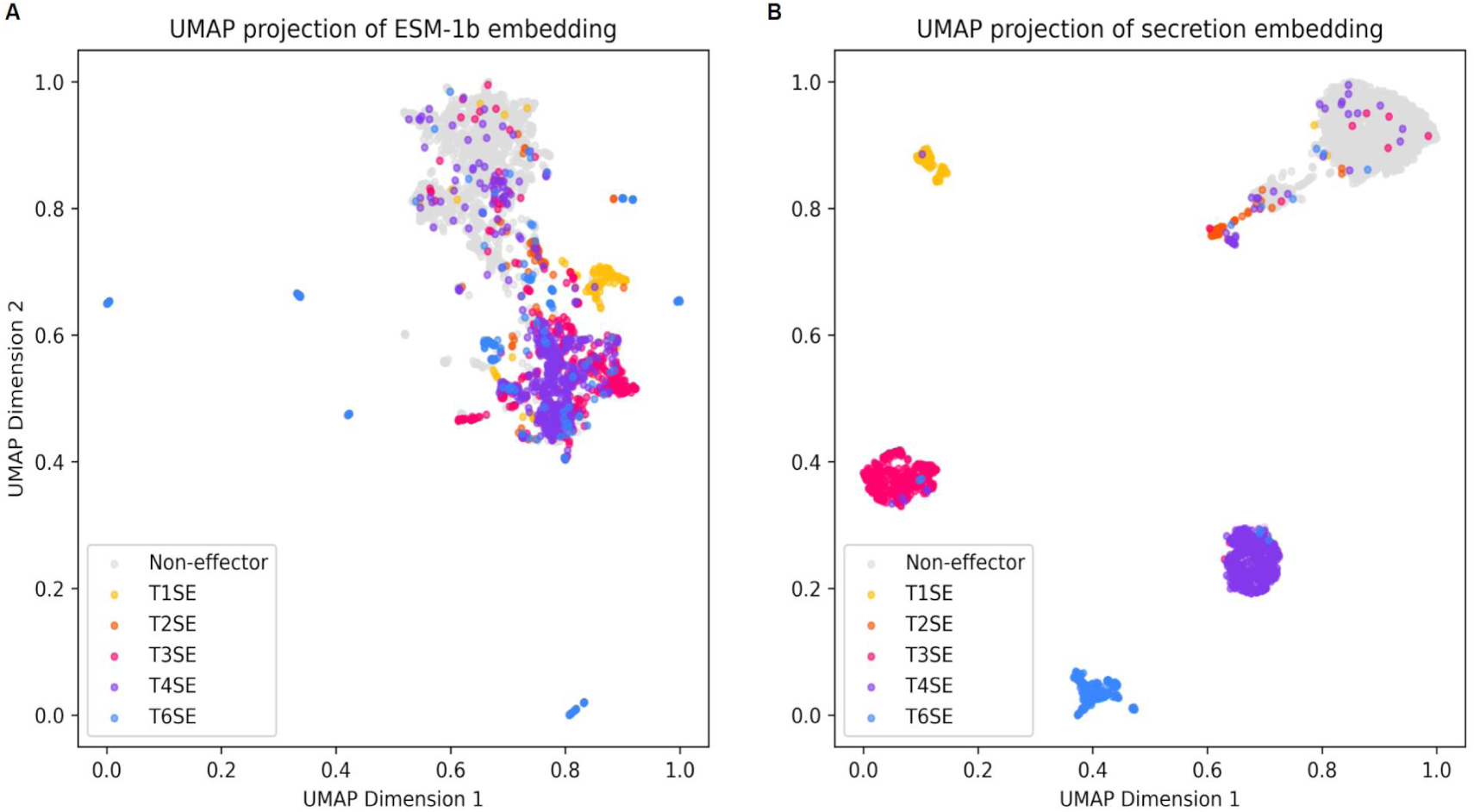
Clustering projection visualization of different types of effector and non-effector proteins. (A) UMAP clustering projections of ESM-1b sequence embeddings. (B) UMAP clustering projections of secretion embeddings from our model.

### Performance evaluation of MoCETSE in effector protein prediction

To comprehensively evaluate the performance and generalization ability of the MoCETSE model, we employed five-fold cross-validation and independent testing. As shown in the Receiver Operating Characteristic (ROC) curves, MoCETSE exhibited strong predictive performance across all categories in the cross-validation experiments (Fig 3A). The Area Under the ROC

**Fig 3.**
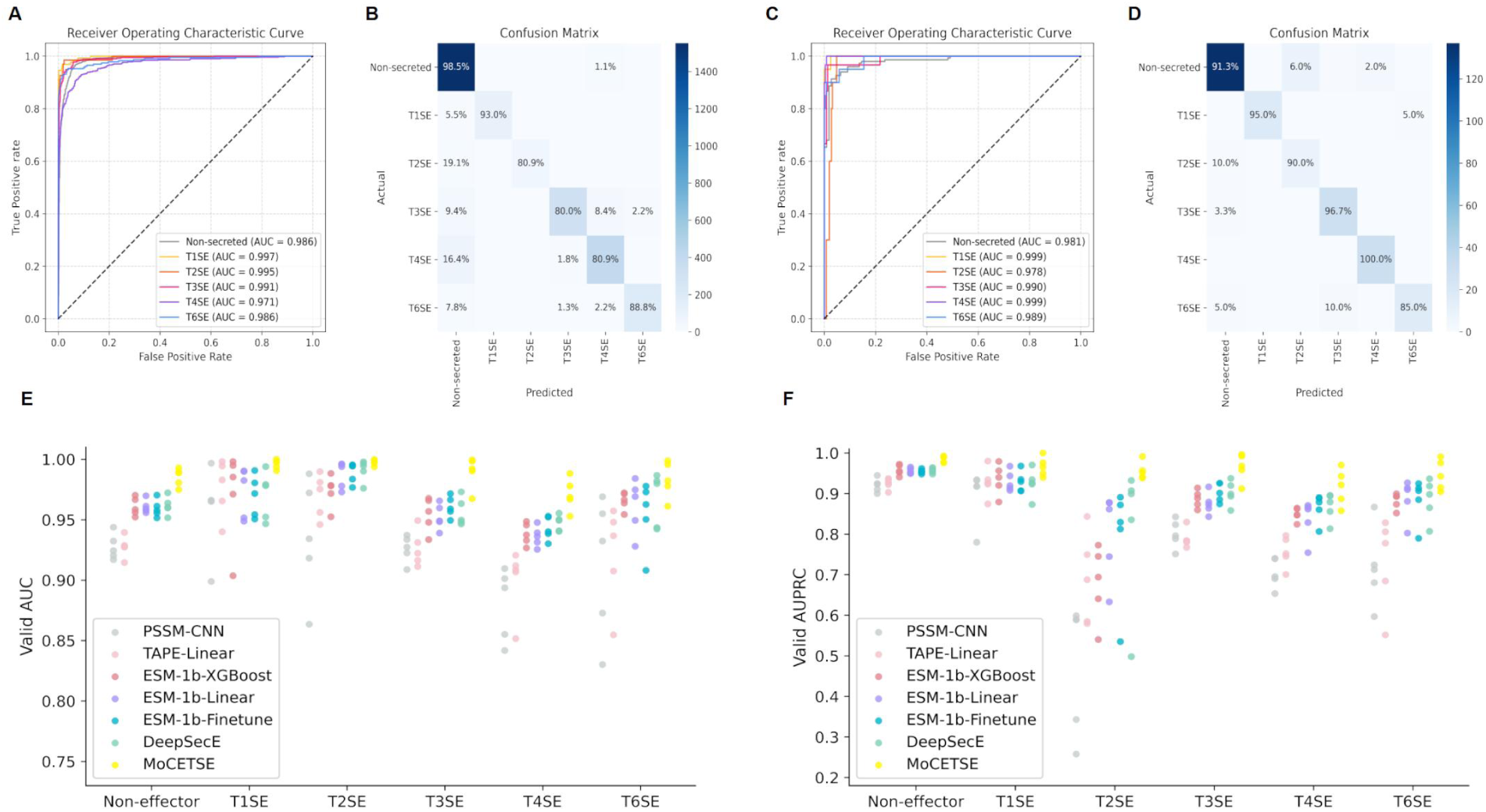
Performance evaluation of MoCETSE for secretion system effector protein prediction. (A to D) Model performance assessed by 5-fold cross-validation (A, B) and independent testing (C and D). ROC curves illustrate the classification of secreted versus non-secreted effectors. Sensitivity values for each class are indicated along the diagonals of the corresponding confusion matrices. (E and F) Comparison of AUC and AUPRC metrics across different model architectures and/or training strategies for the five protein categories under 5-fold cross-validation.

Curve (AUC) values for non-secreted effectors and the five major secreted effectors ranged from 0.971 to 0.997, indicating high classification discriminative ability and achieving a balanced trade-off between sensitivity and specificity. The multi-class confusion matrix provided further insights, illustrating the model’s sensitivity and misclassification rates across all categories (Fig 3B). MoCETSE achieved a classification accuracy of 98.5% for non-secreted effectors, significantly reducing the possibility of false positives. T1SE had the highest sensitivity; due to the scarcity of T2SE data, its sensitivity was relatively low; for T3SE, T4SE, and T6SE, their similar secretion mechanisms involving protein translocation across bacterial membranes may have led to relatively higher misclassification rates. On the independent test set, MoCETSE maintained high generalization performance (Fig 3C). The AUC values for T1SE and T4SE reached 0.999; the AUC for non-effector proteins, T2SE, T3SE, and T6SE were all greater than 0.978, indicating near-perfect discrimination between effector and non-effector proteins. The sensitivities of all categories except T6SE exceeded 90%(Fig 3D). These results highlight the robustness of the model and its potential for reliable application in large-scale and complex effector protein prediction tasks.

### Performance comparison against existing popular models

The MoCETSE model was developed to predict five types of secretion system effector proteins in Gram-negative bacteria, including type I, II, III, IV, and VI effectors. To evaluate its performance, we conducted a comprehensive benchmarking analysis against two existing popular multi-class prediction models, DeepSecE [26] and BastionX [45], as well as nine representative binary classification models. Specifically, T1SEstacker [9] was used for T1SE prediction; Bastion3 [10], T3SEpp [11], and EP3 [12] for T3SE; Bastion4 [13], CNN-T4SE [14], T4SEfinder [15], and T4SEpp [16] for T4SE; and Bastion6 [18] for T6SE. MoCETSE was evaluated on an independent test dataset, and performance metrics were recorded for each effector category. Prediction probabilities for the five effector classes and non-effectors were compared, and statistical analyses were performed to evaluate multi-class classification performance. The evaluation was based on five commonly used metrics: accuracy (ACC), recall (REC), precision (PR), F1 score, and Matthews correlation coefficient (MCC).

To further evaluate the performance and generalization ability of the MoCETSE model in effector protein classification, we conducted extended testing on additional datasets. Comparative analyses against popular models revealed that MoCETSE achieved performance comparable to leading methods for T1SE and T3SE prediction, and outperformed existing approaches for T4SE and T6SE (<u>S1 Table</u>). In the T1SE classification task, MoCETSE achieved comparable performance to DeepSecE, with accuracy (ACC) of 98.8%, F1 score of 0.947, and Matthews correlation coefficient (MCC) of 0.947. It outperformed T1SEstacker, which relies on the amino acid composition of non-RTX C-terminal motifs (ACC: 92.9%, F1: 0.727, MCC: 0.691) (<u>S2A Fig</u>). On the T3SE test set (108 T3SEs and 108 non-effector proteins), MoCETSE achieved an ACC of 91.2%, F1 score of 0.918, and MCC of 0.829. Its performance was closely aligned with that of T3SEpp, a method that integrates multiple biological features (Fig 4A). In the T4SE classification task, evaluated on the CNN-T4SE test set (30 T4SEs and 150 non-secreted effectors), MoCETSE achieved the highest performance among all compared models (ACC: 99.4%, F1: 0.983, MCC: 0.989). In addition, MoCETSE achieved a higher precision (PR) than both DeepSecE and CNN-T4SE, indicating a reduced risk of false optimistic predictions (Fig 4B). For T6SE classification, MoCETSE achieved superior results on the Bastion6 test set (20 T6SEs and 200 non-secreted effectors), with ACC of 98.3%, F1 score of 0.972, and MCC of 0.946 (Fig 4C). Finally, we conducted a further comparison with multiple existing popular models on the independent test dataset of DeepSecE [26], and the results fully confirmed that MoCETSE has the optimal performance (<u>S2 Table</u> and <u>S3 Fig</u>).

**Fig 4.**
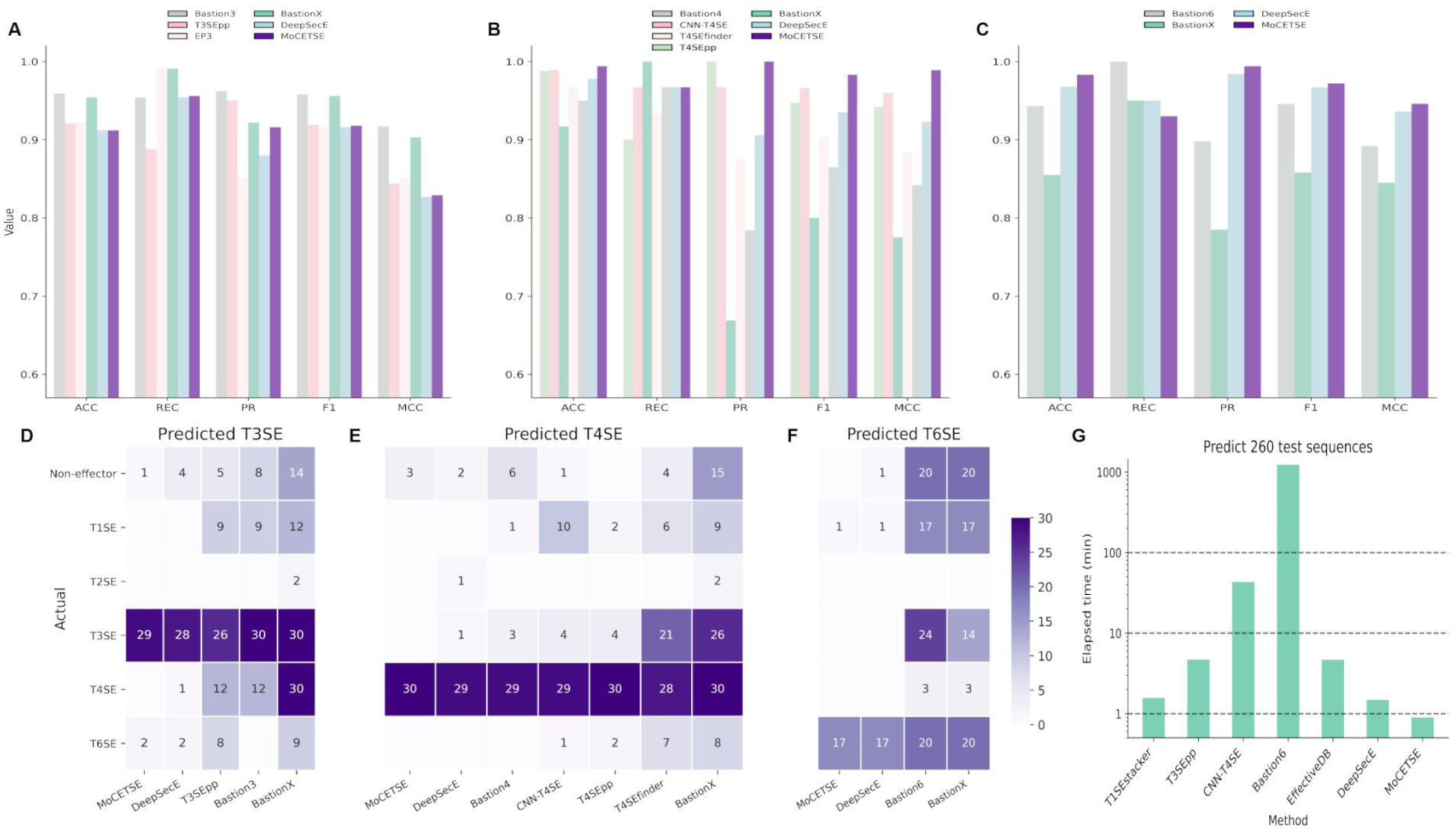
Benchmarking of MoCETSE against existing popular secreted effector protein prediction models. (A to C) Comparisons of MoCETSE with currently popular predictors on T3SE (A), T4SE (B), and T6SE (C). (D to F) Performance evaluation of models based on true positive and false positive identification rates. Heatmaps show the distribution of effector proteins predicted with high confidence. (G) Evaluation of the prediction efficiency of each model on an independent test set containing 260 effector proteins.

Although the benchmark performance of MoCETSE has not yet surpassed Bastion3, the popular T3SE predictor, binary classification methods tend to mispredict other types of secreted effectors as the target type. For example, although Bastion3 can accurately identify all T3SEs in the test set, it misclassifies other types of effectors: 9 T1SEs and 12 T4SEs are incorrectly classified as T3SEs (Fig 4D). In contrast, the MoCETSE model can effectively control the false positive rate when facing multiple effectors, reducing the probability of misjudgment of other types of secreted effectors (Figs 4D to F, <u>S2B Fig</u>). To further evaluate model efficiency, we compared the running time of MoCETSE with other prediction methods in effector classification tasks (Fig 4G). The results showed that MoCETSE completed the prediction in less than 1 minute, and its computational efficiency was significantly improved compared with existing models. The above experimental results fully demonstrate that MoCETSE provides a good balance among accuracy, specificity, and computational efficiency, and its performance is comparable to or even surpasses that of existing popular models.

### Comparison of different foundation methods

To assess the effectiveness of different feature extraction strategies, we constructed two baseline models. The first model employed traditional position-specific scoring matrix (PSSM) features as input, while the second incorporated two advanced pre-trained protein language models, TAPE [46] and ESM-1b [21]. We evaluated the performance of seven models using three metrics: prediction accuracy, F1 score, and area under the precision-recall curve (AUPRC). As shown in Table 1 and Figs 5A-B, the baseline model PSSM-CNN attained an accuracy of 0.799 (95% CI: 0.772–0.826) in five-fold cross-validation, alongside an F1 score of 0.712 (95% CI: 0.649 – 0.774). Incorporating pre-trained protein language models led to clear performance improvements. The TAPEBert (Linear) model attained an accuracy of 0.816 (95% CI: 0.781– 0.851) and an F1 score of 0.764 (95% CI: 0.715–0.813), while the larger-scale ESM-1b (Linear) model further improved accuracy to 0.876 (95% CI: 0.850–0.901) and F1 score to 0.841 (95% CI: 0.795 –0.888). Our proposed MoCETSE model achieved the best overall results across all evaluation metrics, with an accuracy of 0.915 (95% CI: 0.850–0.978) and an F1 score of 0.900 (95% CI: 0.815 – 0.985). Evaluation on an independent test set further confirmed the generalization capacity of MoCETSE; it achieved an accuracy of 0.919 (95% CI: 0.885–0.952) and an F1 score of 0.872 (95% CI: 0.807 – 0.936). The experimental results fully demonstrate that optimizing protein sequence features through ESM-1b and the target preprocessing network can provide effective key feature inputs for the transformer module.

**Fig 5.**
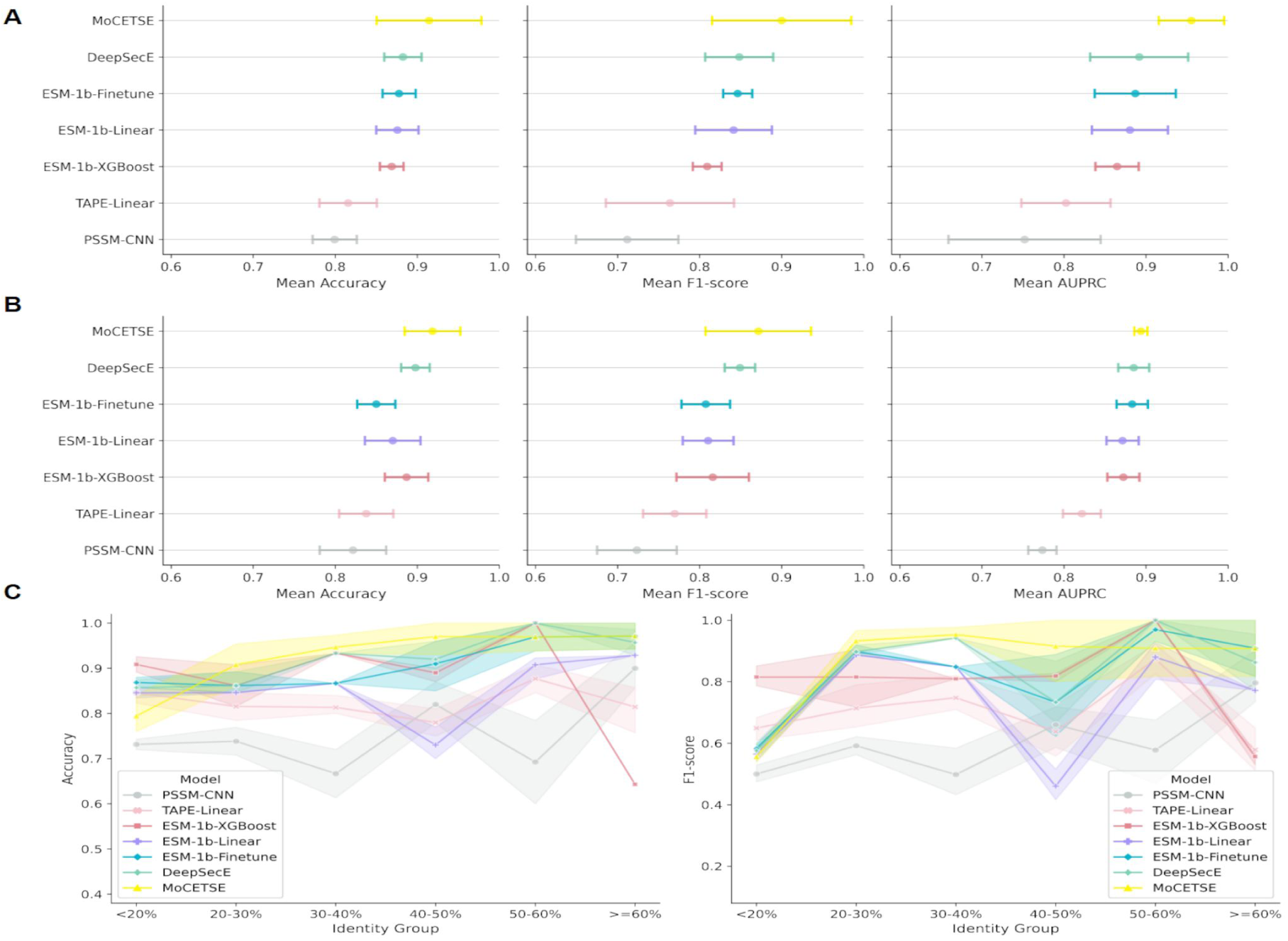
Comparison of model performance between cross-validation and independent test results. (A and B) Average accuracy, F1-score, and AUPRC of different models evaluated by 5-fold cross-validation (A) and independent testing (B), shown with 95% confidence intervals. MoCETSE achieved the highest overall predictive performance. (C) Accuracy (left) and F1-score (right) of each model across sequence identity groups. Shaded regions indicate 95% confidence intervals.

**Table 1.** Model architecture performances across 5-fold cross-validation (CV) and independent (Ind) tests.

| Pre-trained model | Method | ACC |  | F1 |  | AUPRC |  |
| --- | --- | --- | --- | --- | --- | --- | --- |
|  |  | CV | Ind | CV | Ind | CV | Ind |
| None | PSSM-CNN | 0.799 | 0.822 | 0.712 | 0.724 | 0.752 | 0.774 |
| TAPEBert | Linear probing | 0.816 | 0.838 | 0.764 | 0.770 | 0.802 | 0.822 |
| ESM-1b | XGBoost | 0.869 | 0.887 | 0.809 | 0.816 | 0.865 | 0.872 |
| ESM-1b | Linear probing | 0.876 | 0.870 | 0.841 | 0.810 | 0.880 | 0.871 |
| ESM-1b | Fine-tuning | 0.878 | 0.850 | 0.846 | 0.808 | 0.887 | 0.883 |
| ESM-1b | Secretion-specific transformer | 0.883 | 0.898 | 0.848 | 0.849 | 0.892 | 0.885 |
| ESM-1b | Target preprocessing network | **0.915** | **0.919** | **0.900** | **0.872** | **0.955** | **0.894** |

To assess the generalization capability of the model under varying sequence similarity conditions, we stratified the 110 effector proteins in the independent test set into six groups based on sequence identity. The classification performance of MoCETSE and comparison models was then evaluated within each similarity bin (Fig 5C). In terms of AUC, MoCETSE achieved an average improvement of approximately 5.8% over other models, with the largest gains observed in the lowest (<20%) and highest (≥60%) similarity intervals, where AUC increases reached 7.2% and 6.5%, respectively. For F1 score, the average improvement was around 6.3%. Notably, in the high-similarity group (≥60%), MoCETSE outperformed the traditional PSSM-CNN model by 9.1%, and exceeded various ESM-based variants by 4.8% to 6.7%. These results highlight the model’s robustness across diverse sequence similarity ranges and suggest that the integration of pre-trained language representations with targeted preprocessing contributes to more adaptive and reliable feature extraction for effector classification in heterogeneous sequence contexts.

We evaluated the area under the receiver operating characteristic curve (AUC) and area under the precision-recall curve (AUPRC) for non-secreted effectors and five types of secreted effectors in cross-validation(Figs 3E-F). The combined evaluation of AUC and AUPRC can illustrate the model’s ability to distinguish between positive and negative samples and the balance between precision and recall when identifying positive samples. As can be seen from the figures, the use of ESM-1b helps improve the model’s classification performance across effector categories. Compared with other models, MoCETSE achieved the best performance in AUC and AUPRC metrics in most categories, indicating its ability to accurately identify secreted effectors.

### Ablation experiments

To verify the role of each module in the proposed method in improving model performance, we designed a series of ablation experiments. Using the Conv1d-based model as the baseline, we systematically introduced the target preprocessing network (TPN) and relative position encoding (RPE) modules, and evaluated their effects in 5-fold cross-validation (CV) and independent test (Ind) scenarios through two ways: adding them separately and combining them (TPN+RPE). As shown in Table 2 and <u>S4 Fig</u>, the baseline method Conv1d has shown relatively balanced performance in CV and Ind tests, confirming the effectiveness of the basic framework. When the RPE module is introduced alone, the classification results are significantly improved, indicating that the model’s ability to recognize sequence patterns is enhanced; after introducing the TPN module, the model’s performance is further improved, highlighting its potential in optimizing feature representation. However, there are slight fluctuations in some indicators, which show that the advantages of the TPN module need to be coordinated with other mechanisms to be fully released.

**Table 2.** Performance of different methods across 5-fold cross-validation (CV) and independent (Ind) tests in ablation experiments.

| Method | ACC |  | REC |  | PR |  | F1 |  | AUPRC |  |
| --- | --- | --- | --- | --- | --- | --- | --- | --- | --- | --- |
|  | CV | Ind | CV | Ind | CV | Ind | CV | Ind | CV | Ind |
| Conv1d | 0.883 | 0.898 | 0.823 | 0.882 | 0.882 | 0.834 | 0.848 | 0.849 | 0.892 | 0.885 |
| Conv1d+RPE | 0.902 | 0.864 | 0.871 | 0.867 | 0.915 | 0.786 | 0.898 | 0.812 | 0.951 | 0.872 |
| TPN | 0.911 | 0.875 | 0.869 | 0.858 | 0.927 | 0.808 | 0.897 | 0.820 | 0.954 | 0.880 |
| TPN+RPE | **0.915** | **0.919** | **0.873** | **0.895** | **0.937** | **0.863** | **0.900** | **0.872** | **0.955** | **0.894** |

It is worth noting that after the fusion of TPN and RPE dual modules, the model performance has achieved a breakthrough improvement. This result reveals the functional complementarity of the two modules: RPE strengthens the accuracy of sequence feature extraction, while TPN plays a key role in feature interaction and decision optimization. The two work together to build a more efficient classification pathway. The comparison of ablation experiments shows that the dual-module fusion is not a simple superposition of performance, but through the collaboration at the mechanism level, it breaks through the performance bottleneck of a single module, providing an effective solution for the model to adapt to complex feature scenarios in the effector protein sequence classification task.

### Relative position attention identification sequence motifs

Signal peptides are key identifiers of secreted effectors and are generally located at the N-terminus or C-terminus of the sequence. In the study of the Dot/Icm T4SS secreted protein VpdB [7] in *Legionella pneumophila* Philadelphia 1 strain (UniProt accession number Q5ZW60), the MoCETSE model introduced a relative position multi-head attention mechanism to accurately capture the relative distance relationships between amino acids in the sequence, thereby effectively mining functional signals generated by spatial proximity and structural association. Experimental results showed that MoCETSE successfully identified the N-terminal sequence consistent with type Ⅳ secretion signal characteristics and detected the C-terminal region predicted to be related to the secretion process (Fig 6).

**Fig 6.**
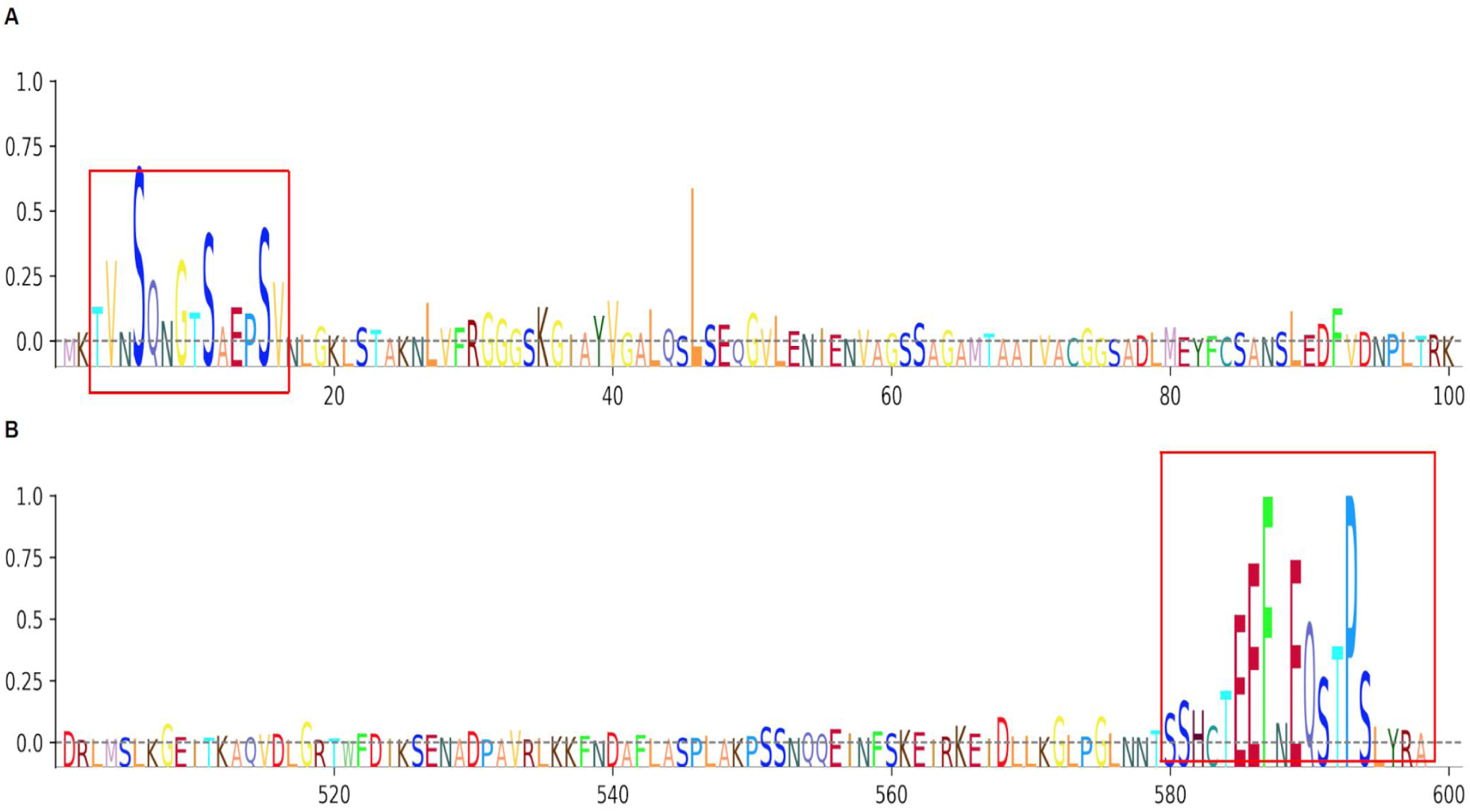
Visualization of sequence features in the Dot/Icm T4SS effector VpdB identified by MoCETSE. (A and B) Visualization of the N-terminal (A) and C-terminal (B) regions of VpdB. The N-terminus displays features consistent with known type IV secretion signals, while the C-terminus reveals sequence patterns potentially implicated in the secretion process.

## Discussion

This study proposed MoCETSE, a deep learning framework that advances the prediction and classification of bacterial secretion system effector proteins by balancing accuracy, interpretability, and computational efficiency. MoCETSE uses a pre-trained protein language model for sequence encoding; through the innovative module target preprocessing network for dimensionality reduction and feature extraction, it significantly improves the model’s prediction accuracy while reducing redundant noise and improving computational efficiency (Fig 1B); the combination of relative position encoding and multi-head attention further enables the model to capture long-range dependencies and highlight sequence motifs related to effector proteins, providing valuable biological interpretability while delivering high predictive performance (Fig 1C).

MoCETSE established a new benchmark for multi-class effector protein classification by outperforming or matching existing popular methods across different effector protein tasks (Figs 4A to C). Benchmarking showed that MoCETSE not only outperformed the binary classifier CNN-T4SE and the multi-class classifier DeepSecE, but also exhibited a lower false positive rate and superior specificity in scenarios involving mixed effector proteins (Figs 4D to F). MoCETSE achieved the best performance in T4SE and T6SE classification tasks, while maintaining competitiveness in T1SE and T3SE, highlighting the strong predictive ability of the model. In addition, compared with existing predictors, MoCETSE’s efficient framework produced faster inference, making it more suitable for large-scale or real-time applications (Fig 4G).

MoCETSE demonstrated a strong ability to generalize across different sequence contexts. To further evaluate this ability, we stratified the independent test set according to sequence similarity and observed that MoCETSE consistently maintained high accuracy across different homology ranges (Fig 5C). These findings indicate that the model is robust to sequence heterogeneity and demonstrates higher reliability when encountering previously uncharacterized or distantly related effector proteins.

MoCETSE effectively improves the prediction performance of bacterial secretion system effector proteins through its innovative architectural design. The target preprocessing network module uses parallel convolutional experts to capture multi-scale sequence signals that traditional architectures might overlook, while suppressing noise and enhancing the model’s ability to recognize complex effector protein-specific patterns. At the same time, the multi-head attention mechanism integrated with relative position encoding enables MoCETSE to focus on biologically meaningful regions, such as the N- and C-termini, providing interpretable outputs consistent with known secretion-associated motifs (Fig 6).

Although the MoCETSE model has achieved excellent performance in predicting bacterial secretion system effector proteins, it still has certain limitations. Currently, the available datasets are relatively limited and may not fully cover the complete features of natural effector proteins. Additionally, there is an imbalance in the distribution among different classes—for example, the number of trainable samples for T2SE is relatively small. This may weaken the model’s discriminative ability for low-frequency classes and limit its generalization performance. Future work will focus on expanding the dataset by incorporating a broader range of bacterial species and newly discovered effector proteins to enhance the model’s generalization capability and robustness. Additionally, we may integrate larger-scale or more advanced pre-trained protein language models (such as SaProt and ESM3) [47,48] to obtain higher-quality and biologically more meaningful feature embeddings, thereby improving the prediction accuracy of downstream tasks (secreted effectors). However, this requires balancing both model interpretability and computational resource availability.

## Conclusion

MoCETSE effectively improves the prediction accuracy and generalization ability of the model by integrating amino acid sequence features, evolutionary information provided by ESM-1b, multi-scale representations extracted by the mixture-of-convolution experts, and secretion signals learned by the transformer. This approach facilitates the initial identification of effector proteins and enhances comprehensive insights into viral pathogenic mechanisms. MoCETSE highlights how to combine large-scale language models with biological information modules to create a more powerful framework for researching pathogen-related proteins. With the continuous development of large language models, future research will focus on adopting more advanced and larger-scale protein language models to enhance the robustness and practicality of MoCETSE, thereby further expanding its capabilities in handling higher-dimensional and more complex biological tasks.

## Acknowledgments

We thank Jiani Chen and Zhouying Li for discussion on related topics.

## Supporting information

S1 Table. Performance comparison of the proposed method versus existing popular approaches on benchmark datasets for predicting T1SE, T3SE, T4SE, and T6SE.

(PDF)

S2 Table. Performance comparison of our method versus existing popular approaches on DeepSecE benchmark datasets for T1SE, T3SE, T4SE, and T6SE prediction.

(PDF)

S1 Fig. Protein sequence length information in the training set and test set. (A–B) Distribution histograms illustrating the composition of protein sequence lengths in the training dataset (A) and the test dataset (B). The x-axis denotes protein sequence length (in amino acids), while the y-axis indicates the frequency of proteins within each length interval.

(PDF)

S2 Fig. The performance of MoCETSE was systematically compared with that of existing popular models in the task of predicting type I secreted effectors (T1SEs). (A) Model performance was evaluated on a test set comprising 150 non-effector proteins and 20 T1SEs, using metrics including accuracy (ACC), recall (REC), precision (PR), F1-score, and Matthews correlation coefficient (MCC). (B) True positive and false positive predictions for T1SEs were compared across different models. A heatmap illustrates the distribution of proteins predicted as T1SEs across various effector classes by each model.

(PDF)

S3 Fig. MoCETSE was evaluated against several existing popular models on the test set provided by DeepSecE. (A–C) show the performance of different models on the prediction tasks for T3SE (A), T4SE (B), and T6SE (C), respectively, across multiple evaluation metrics.

(PDF)

S4 Fig. Performance comparison of different module configurations (Conv1d, Conv1d+RPE, TPN, TPN+RPE) in ablation experiments. (A) Results from 5-fold cross-validation. (B) Results on the independent test set.

(PDF)

## References

1. Costa TR, Felisberto-Rodrigues C, Meir A, Prevost MS, Redzej A, Trokter M, et al. Secretion systems in Gram-negative bacteria: Structural and mechanistic insights. Nat Rev Microbiol. 2015;13(6):343–359. 10.1038/nrmicro3456 PMID: 25978706

2. Green ER, Mecsas J. Bacterial secretion systems: An overview. Microbiol Spectr. 2016;4(1): 10.1128/microbiolspec.VMBF-0012-2015 PMID: 26999395

3. Hui X, Chen Z, Zhang J, Lu M, Cai X, Deng Y, et al. Computational prediction of secreted proteins in gram-negative bacteria. Comput Struct Biotechnol J. 2021;19:1806–1828. 10.1016/j.csbj.2021.03.019 PMID: 33897982

4. Sanchez-Garrido J, Ruano-Gallego D, Choudhary JS, Frankel G. The type III secretion system effector network hypothesis. Trends Microbiol. 2022;30(6):524–533. 10.1016/j.tim.2021.10.007 PMID: 34840074

5. Singh RP, Kumari K. Bacterial type VI secretion system (T6SS): an evolved molecular weapon with diverse functionality. Biotechnol Lett. 2023;45(3):309–331. 10.1007/s10529-023-03354-2 PMID: 36683130

6. Ruano-Gallego D, Sanchez-Garrido J, Kozik Z, Núñez-Berrueco E, Cepeda-Molero M, Mullineaux-Sanders C, et al. Type III secretion system effectors form robust and flexible intracellular virulence networks. Science. 2021;371(6534):eabc9531. 10.1126/science.abc9531 PMID: 33707240

7. Böck D, Hüsler D, Steiner B, Medeiros JM, Welin A, Radomska KA, et al. The Polar Legionella Icm/Dot T4SS Establishes Distinct Contact Sites with the Pathogen Vacuole Membrane. mBio. 2021;12(5):e0218021. 10.1128/mBio.02180-21 PMID: 34634944

8. Colautti J, Kelly SD, Whitney JC. Specialized killing across the domains of life by the type VI secretion systems of Pseudomonas aeruginosa. Biochem J. 2025;482(1):1–15. 10.1042/BCJ20230240 PMID: 39774785

9. Chen Z, Zhao Z, Hui X, Zhang J, Hu Y, Chen R, et al. T1SEstacker: A Tri-Layer Stacking Model Effectively Predicts Bacterial Type 1 Secreted Proteins Based on C-Terminal Non-repeats-in-Toxin-Motif Sequence Features. Front Microbiol. 2022;12:813094. 10.3389/fmicb.2021.813094 PMID: 35211101

10. Wang J, Li J, Yang B, Xie R, Marquez-Lago TT, Leier A, et al. Bastion3: a two-layer ensemble predictor of type III secreted effectors. Bioinformatics. 2019;35(12):2017–2028. 10.1093/bioinformatics/bty914 PMID: 30388198

11. Hui X, Chen Z, Lin M, Zhang J, Hu Y, Zeng Y, et al. T3SEpp: an Integrated Prediction Pipeline for Bacterial Type III Secreted Effectors. mSystems. 2020;5(4):e00288–20. 10.1128/mSystems.00288-20 PMID: 32753503

12. Li J, Wei L, Guo F, Zou Q. EP3: an ensemble predictor that accurately identifies type III secreted effectors. Brief Bioinform. 2021;22(2):1918–1928. 10.1093/bib/bbaa008 PMID: 32043137

13. Wang J, Yang B, An Y, Marquez-Lago T, Leier A, Wilksch J, et al. Systematic analysis and prediction of type IV secreted effector proteins by machine learning approaches. Brief Bioinform. 2019;20(3):931–951. 10.1093/bib/bbx164 PMID: 29186295

14. Hong J, Luo Y, Mou M, Fu J, Zhang Y, Xue W, et al. Convolutional neural network-based annotation of bacterial type IV secretion system effectors with enhanced accuracy and reduced false discovery. Brief Bioinform. 2020;21(5):1825–1836. 10.1093/bib/bbz120 PMID: 31860715

15. Zhang Y, Zhang Y, Xiong Y, Wang H, Deng Z, Song J, et al. T4SEfinder: a bioinformatics tool for genome-scale prediction of bacterial type IV secreted effectors using pre-trained protein language model. Brief Bioinform. 2022;23(1):bbab420. 10.1093/bib/bbab420 PMID: 34657153

16. Hu Y, Wang Y, Hu X, Chao H, Li S, Ni Q, et al. T4SEpp: A pipeline integrating protein language models to predict bacterial type IV secreted effectors. Comput Struct Biotechnol J. 2024;23:801–812. 10.1016/j.csbj.2024.01.015 PMID: 38328004

17. Li J, He S, Zhang J, Zhang F, Zou Q, Ni F. T4Seeker: a hybrid model for type IV secretion effectors identification. BMC Biol. 2024;22(1):259. 10.1186/s12915-024-02064-z PMID: 39543674

18. Wang J, Yang B, Leier A, Marquez-Lago TT, Hayashida M, Rocker A, et al. Bastion6: a bioinformatics approach for accurate prediction of type VI secreted effectors. Bioinformatics. 2018;34(15):2546–2555. 10.1093/bioinformatics/bty155 PMID: 29547915

19. Sen R, Nayak L, De RK. PyPredT6: A python-based prediction tool for identification of Type VI effector proteins. J Bioinform Comput Biol. 2019;17(3):1950019. 10.1142/S0219720019500197 PMID: 31288641

20. Lin Z, Akin H, Rao R, Hie B, Zhu Z, Lu W, et al. Evolutionary-scale prediction of atomic-level protein structure with a language model. Science. 2023;379(6637):1123-1130. 10.1126/science.ade2574 PMID: 36927031

21. Rives A, Meier J, Sercu T, Goyal S, Lin Z, Liu J, et al. Biological structure and function emerge from scaling unsupervised learning to 250 million protein sequences. Proc Natl Acad Sci U S A. 2021;118(15):e2016239118. 10.1073/pnas.2016239118 PMID: 33876751

22. Rao R, Meier J, Sercu T, Ovchinnikov S, Rives A, Jones P, et al. Transformer protein language models are unsupervised structure learners. In: Proceedings of the 9th International Conference on Learning Representations; 2021 May 3 – 7; Vienna, Austria. Vienna (Austria): ICLR Press; 2021. p. 123–130.

23. Vaswani A, Shazeer N, Parmar N, Uszkoreit J, Jones L, Gomez A, et al. Attention is all you need. In: Proceedings of the 31st International Conference on Neural Information Processing Systems (NIPS 2017); 2017 Dec 4 – 9; Long Beach, CA. New York: Curran Associates; 2017. p. 5998–6008.

24. Brandes N, Ofer D, Peleg Y, Rappoport N, Linial M. ProteinBERT: a universal deep-learning model of protein sequence and function. Bioinformatics. 2022;38(8):2102–2110. 10.1093/bioinformatics/btac020 PMID: 35020807

25. Chen JY, Wang JF, Hu Y, Li XH, Qian YR, Song CL. Evaluating the advancements in protein language models for encoding strategies in protein function prediction: a comprehensive review. Front Bioeng Biotechnol. 2025;13:1506508. 10.3389/fbioe.2025.1506508 PMID: 39906415

26. Zhang Y, Guan J, Li C, Wang Z, Deng Z, Gasser RB, et al. DeepSecE: A Deep-Learning-Based Framework for Multiclass Prediction of Secreted Proteins in Gram-Negative Bacteria. Research (Wash D C). 2023;6:0258. 10.34133/research.0258 PMID: 37886621

27. Liao XJ, He TT, Liu LY, Jiang XL, Sun SS, Deng YH, et al. Unraveling and characterization of novel T3SS effectors in Edwardsiella piscicida. mSphere. 2023;8(5):e0034623. 10.1128/msphere.00346-23 PMID: 37642418

28. Hendrycks D, Gimpel K. Gaussian error linear units (GELUs) [Preprint]. arXiv:1606.08415 [cs.LG]. 2016 Jun 27 [revised 2023 Jun 6; version 5]. Available from: https://arxiv.org/abs/1606.08415

29. Madani A, Krause B, Greene ER, Subramanian S, Mohr BP, Holton JM, et al. Large language models generate functional protein sequences across diverse families. Nat Biotechnol. 2023;41(8):1099–1106. 10.1038/s41587-022-01618-2 PMID: 36702895

30. Teufel F, Almagro Armenteros JJ, Johansen AR, Gíslason MH, Pihl SI, Tsirigos KD, et al. SignalP 6.0 predicts all five types of signal peptides using protein language models. Nat Biotechnol. 2022;40(7):1023–1025. 10.1038/s41587-021-01156-3 PMID: 34980915

31. Ahmad S, Jose da Costa Gonzales L, Bowler-Barnett EH, Rice DL, Kim M, Wijerathne S, et al. The UniProt website API: facilitating programmatic access to protein knowledge. Nucleic Acids Res. 2025;53(W1):W547–W553. 10.1093/nar/gkaf394 PMID: 40331428

32. Jacobs RA, Jordan MI, Nowlan SJ, Hinton GE. Adaptive Mixtures of Local Experts. Neural Comput. 1991;3(1):79–87. 10.1162/neco.1991.3.1.79 PMID: 31141872

33. Dai D, Deng C, Zhao C, Xu RX, Gao H, Chen D, et al. DeepSeekMoE: towards ultimate expert specialization in mixture-of-experts language models [Preprint]. arXiv:2401.05639 [cs.CL]. 2024 Jan 11. Available from: https://arxiv.org/abs/2401.05639

34. Huang Q, An Z, Zhuang N, Tao M, Zhang C, Jin Y, et al. Harder tasks need more experts: dynamic routing in MoE models [Preprint]. arXiv:2403.07679 [cs.LG]. 2024 Mar 12. Available from: https://arxiv.org/abs/2403.07679

35. Liu H, Xia M, Gao T, Wang R, Chen D. Gating Dropout: Communication-efficient Regularization for Sparsely Activated Transformers. arXiv [Preprint]. 2022 May 27. Available from: https://arxiv.org/abs/2205.14336

36. Zhang Y, Cai R, Chen T, Zhang G, Zhang H, Chen P, et al. Robust mixture-of-expert training for convolutional neural networks [Preprint]. arXiv:2308.09751 [cs.CV]. 2023 Aug 19. Available from: https://arxiv.org/abs/2308.09751

37. Cho K, van Merrienboer B, Gulcehre C, Bahdanau D, Bougares F, Schwenk H, et al. Learning phrase representations using RNN encoder-decoder for statistical machine translation [Preprint]. arXiv:1406.1078 [cs.CL]. 2014 Jun 3 [revised 2014 Sep 3; version 3]. Available from: https://arxiv.org/abs/1406.1078

38. Gehring J, Auli M, Grangier D, Yarats D, Dauphin YN. Convolutional sequence to sequence learning [Preprint]. arXiv:1705.03122 [cs.CL]. 2017 May 8 [revised 2017 Jul 25; version 3]. Available from: https://arxiv.org/abs/1705.03122

39. Zhou Y, Zhao X, Guo X, Li J, Liu S. Design of a modified transformer architecture based on relative position coding. Int J Comput Intell Syst. 2021;14(1):89–99. 10.1007/s44196-023-00345-z

40. Shaw P, Uszkoreit J, Vaswani A. Self-attention with relative position representations. In: Proceedings of the 2018 Conference on Neural Information Processing Systems (NeurIPS 2018); 2018 Dec 2 – 8; Montréal, Canada. Curran Associates; 2018. p. 4644 – 4652. 10.18653/v1/N18-2074

41. Zhu H, Hao H, Yu L. Identifying disease-related microbes based on multi-scale variational graph autoencoder embedding Wasserstein distance. BMC Biol. 2023;21(1):294. 10.1186/s12915-023-01796-8 PMID: 38115088

42. McDermott MBA, Zhang H, Hansen LH, Angelotti G, Gallifant J, et al. A closer look at AUROC and AUPRC under class imbalance [Preprint]. arXiv:2401.05706 [cs.LG]. 2024 Jan 11 [revised 2025 Jan 13; version 4]. Available from: https://arxiv.org/abs/2401.05706

43. Tareen A, Kinney JB. Logomaker: beautiful sequence logos in Python. Bioinformatics. 2020;36(7):2272–2274. 10.1093/bioinformatics/btz921 PMID: 31821414

44. McInnes L, Healy J, Melville J. UMAP: uniform manifold approximation and projection for dimension reduction [Preprint]. arXiv:1802.03426 [cs.LG]. 2018 Feb 9 [revised 2020 Sep 18; version 3]. Available from: https://arxiv.org/abs/1802.03426

45. Wang J, Li J, Hou Y, Dai W, Xie R, Marquez-Lago TT, et al. BastionHub: a universal platform for integrating and analyzing substrates secreted by Gram-negative bacteria. Nucleic Acids Res. 2021;49(D1):D651–D659. 10.1093/nar/gkaa899 PMID: 33084862

46. Rao R, Bhattacharya N, Thomas N, Duan Y, Chen X, Canny J, et al. Evaluating Protein Transfer Learning with TAPE. Adv Neural Inf Process Syst. 2019;32:9689–9701. 10.48550/arXiv.1906.08230 PMID: 33390682

47. Su J, Han C, Zhou Y, Shan J, Zhou X, Yuan F. SaProt: Protein language modeling with structure-aware vocabulary. In: Proceedings of the International Conference on Learning Representations (ICLR 2024); 2024 May; Kigali, Rwanda. ICLR Press; 2024. Poster Spotlight.

48. Hayes T, Rao R, Akin H, Sofroniew NJ, Oktay D, Lin Z, et al. Simulating 500 million years of evolution with a language model. Science. 2025;387(6736):850-858. 10.1126/science.ads0018 PMID: 39818825

